# Dosage sensitivity of X-linked genes in human embryonic single cells

**DOI:** 10.1101/291724

**Authors:** Jian-Rong Yang, Xiaoshu Chen

**Author notes:** Correspondence to: Xiaoshu Chen, Department of Medical Genetics, Zhongshan School of Medicine, Sun Yat-sen University, 1212 Medical Science and Technology Building, 74 Zhongshan 2^nd^ Road, Guangzhou, Guangdong 510080. Jian-Rong Yang, Department of Biology, Zhongshan School of Medicine, Sun Yat-sen University, 1227 Medical Science and Technology Building, 74 Zhongshan 2nd Road, Guangzhou, Guangdong 510080, OR.

## Abstract

Fifty years ago, Susumu Ohno proposed that the expression levels of X-linked genes have doubled as dosage compensation for autosomal genes due to degeneration of Y-linked homologs during evolution of mammalian sex chromosomes. Recent studies have nevertheless shown that the X to autosome expression ratio equals ~1 in haploid human parthenogenetic embryonic stem (pES) cells and ~0.5 in diploid pES cells, thus refuting Ohno’s hypothesis. Here, by reanalyzing a RNA-seq-based single-cell transcriptome dataset of human embryos (Petropoulos, et al. 2016), we found that from the 8-cell stage until the time-point just prior to implantation, the expression levels of X-linked genes are not two-fold upregulated in male cells and gradually decrease from two-fold in female cells. This observation suggests that the expression levels of X-linked genes are imbalanced, with autosomal genes starting from the early 8-cell stage, and that the dosage conversion is fast, such that the X:AA expression ratio reaches ~0.5 in no more than a week. Additional analyses of gene expression noise further suggest that the dosage sensitivity of X-linked genes is weaker than that of autosomal genes in differentiated female cells, which contradicts a key assumption of Ohno’s hypothesis. Moreover, the dosage-sensitive housekeeping genes are preferentially located on autosomes, implying selection against X-linkage for dosage-sensitive genes. Our results collectively suggest an alternative to Ohno’s hypothesis that X-linked genes are less likely to be dosage sensitive than autosomal genes.

## Introduction

Mammalian sex chromosomes evolved from a pair of autosomes, in which the evolutionary degeneration of Y potentially causes dosage imbalance between X-linked and autosomal genes. Fifty years ago, Ohno proposed that the per-allele expression levels of X-linked genes should be doubled to re-balance the dosage (Ohno 1967) in both male cells, in which only one X chromosome exists, and female cells, in which one of the two X chromosomes is inactivated (Goto and Monk 1998). Ohno’s hypothesis formed the theoretical foundation for the current model of mammalian sex chromosome evolution and sex chromosome dosage compensation (Charlesworth 1996; Payer and Lee 2008). In 2006, a set of microarray-based gene expression profiles in human somatic tissues facilitated the first genome-wide empirical test of Ohno’s hypothesis (Nguyen and Disteche 2006). It was found that the gene expression ratio between one active X (X_a_) and two autosomes (AA) is approximately 1, or X_a_:AA ~1, lending support to Ohno’s hypothesis (Nguyen and Disteche 2006).

However, gene expressions are reflected by probe-specific affinities in microarrays, which perform poorly in quantifying between-chromosome expression ratios (Xiong, et al. 2010). Ohno’s hypothesis was thus re-examined using RNA-Seq-based expression profiles (Xiong, et al. 2010) and proteomic data (Chen and Zhang 2015), and X_a_:AA was found to be ~0.5. Additionally, comparison between human X-linked genes and proto-X genes (i.e., the autosomal progenitors of the X-linked genes) suggested no change in per-allele expression levels during mammalian X chromosome evolution (Julien, et al. 2012; Lin, et al. 2012). Furthermore, the X to autosome expression ratio (X_a_:A) in human parthenogenetic embryonic stem (pES) haploid cells (containing one active X and one set of autosomes) was found to be ~1 (Chen and Zhang 2016; Nguyen and Disteche 2006). Intriguingly, X-linked genes encoding components of large protein complexes, which are supposed to be dosage-sensitive, are upregulated in both diploid (Lin, et al. 2012; Pessia, et al. 2012) and haploid cells (Chen and Zhang 2016), leading to dosage imbalance in haploid cells. More recently, X-linked genes are found upregulated in germ cells but not in soma cells (Sangrithi, et al. 2017). Collectively, these results have largely refuted the universality of Ohno’s theory (Mank, 2013), whereas its scope of application remains debated. In contrast, an alternative scenario emerges, i.e., X-linked genes are insensitive to the two-fold expression change caused by either evolutionary degeneration of the Y-linked homologs, or the physiological transition of ploidy (as in meiosis and zygote formation) or X-inactivation. In this study, we examined this “insensitive X hypothesis”.

We reasoned that a gene with higher dosage sensitivity should display lower expression variance. Similar logic has been invoked in previous studies, in which genes with lower expression variance between individuals are considered under stronger selection on the dosage of expression (Mullon, et al. 2015). Instead of estimating expression variance between biological replicates (Mullon, et al. 2015; Yin, et al. 2009), we took advantage of a recently published single-cell RNA-seq study of human embryos (Petropoulos, et al. 2016) to directly gauge the level of expression noise for each gene (Newman, et al. 2006). This dataset includes the transcriptomes of 1,529 individual cells at embryonic days (E) 3–7 from 88 human preimplantation embryos, with a temporal span from the 8-cell stage up to the time-point just prior to implantation (Petropoulos, et al. 2016). There are a total of 15,633 genes expressed in at least 5 sequenced cells with RPKM (***R***eads ***P***er ***K***ilobase exon model and per ***M***illion mapped reads) no less than 10. This dataset has allowed determination of the sex of each cell by the expression of Y-linked genes and categorization of individual cells into three clearly segregating lineages, namely, trophectoderm (TE), primitive endoderm (PE), and epiblast (EPI) lineages. Analyses of this dataset serendipitously revealed biallelic transcription of *XIST* throughout the progression of X expression dampening, and X-linked genes are transcribed from both alleles in the female preimplantation embryo (Petropoulos, et al. 2016). This phenomenon is in contrast to the complete silencing of one randomly selected X chromosome in later development (Petropoulos, et al. 2016).

The single-cell transcriptome data of preimplantation embryos gives us a unique opportunity to test key predictions of the insensitive X hypothesis. First, during the physiological process of X inactivation in female cells, the dosage balance between sex chromosomes and autosomes is expected to change, which should remain unchanged according to Ohno’s theory. Second, the dosage sensitivity, as reflected by diminished expression variation among individual cells, should be lower for X-linked genes than for autosomal genes. Third, the expression of X-linked genes should be more variable than well-defined dosage-sensitive genes, such as housekeeping genes. In the following sections, we individually test these predictions.

### Expression levels of X-linked genes are imbalanced with autosomal genes from the early 8-cell stage

As Ohno’s hypothesis concerns genes that existed before the origin of mammalian X, we evaluated previous studies (Xiong, et al. 2010;Lin, et al. 2012) that focused on human genes with one-to-one orthologs in chicken. For a fair comparison of the expression levels, we need to choose unbiased sets of X-lined and autosomal genes. Two strategies were previously employed to that end. On the one hand, a single lower limit was used to choose active genes on both X-linked and autosomal genes. One the other hand, identical fractions of highly expressed genes were chosen from X and autosomes. Mathematically, the former strategy is only appropriate if X:AA ≈ 1, but overestimates the ratio when X:AA < 1. The latter strategy, however, gives unbiased estimation of X:AA regardless the real ratio (He, et al. 2011). We thus compared, for each day and each lineage, the fraction of X-linked genes whose mean expression level in all single cells is no less than 10 RPKM (Petropoulos, et al. 2016) and the same fraction of autosomal genes with the highest expression levels (Chen and Zhang 2016). The ratio of median mRNA expression levels between X-link genes and autosomal genes was then calculated and referred to as the X:AA expression ratio.

We found that the X:AA expression ratio in male cells is ~0.5 regardless of lineage and time point (triangles in **Fig. 1a**). Specifically, the 90% confidence interval of the estimated X:AA expression ratio overlaps with 0.5 but not 1 (**Fig. 1a**). This result is similar to a previous observation made by RNA-seq in human male diploid cells (Xiong, et al. 2010). On the other hand, the X:AA expression ratio of female cells gradually decreases from ~0.75 at E3, to ~0.5 at E7 (circles in **Fig. 1a**). It is also noteworthy that the 90% confidence interval of the X:AA expression ratio of female cells never reaches the prediction made in Ohno’s hypothesis. This result is observed in every lineage (**Fig. 1a**) and remains the same even when we use a less stringent cut-off (RPKM ≥5, **Fig. S1a**) when filtering expressed X-linked genes. The slightly higher X:AA expression ratio in early time points is likely a transition state between haploid cells (X:A ~ 1) and diploid cells (X:AA ~ 0.5) as the maternal to zygotic transition occurs.

**Figure 1.**
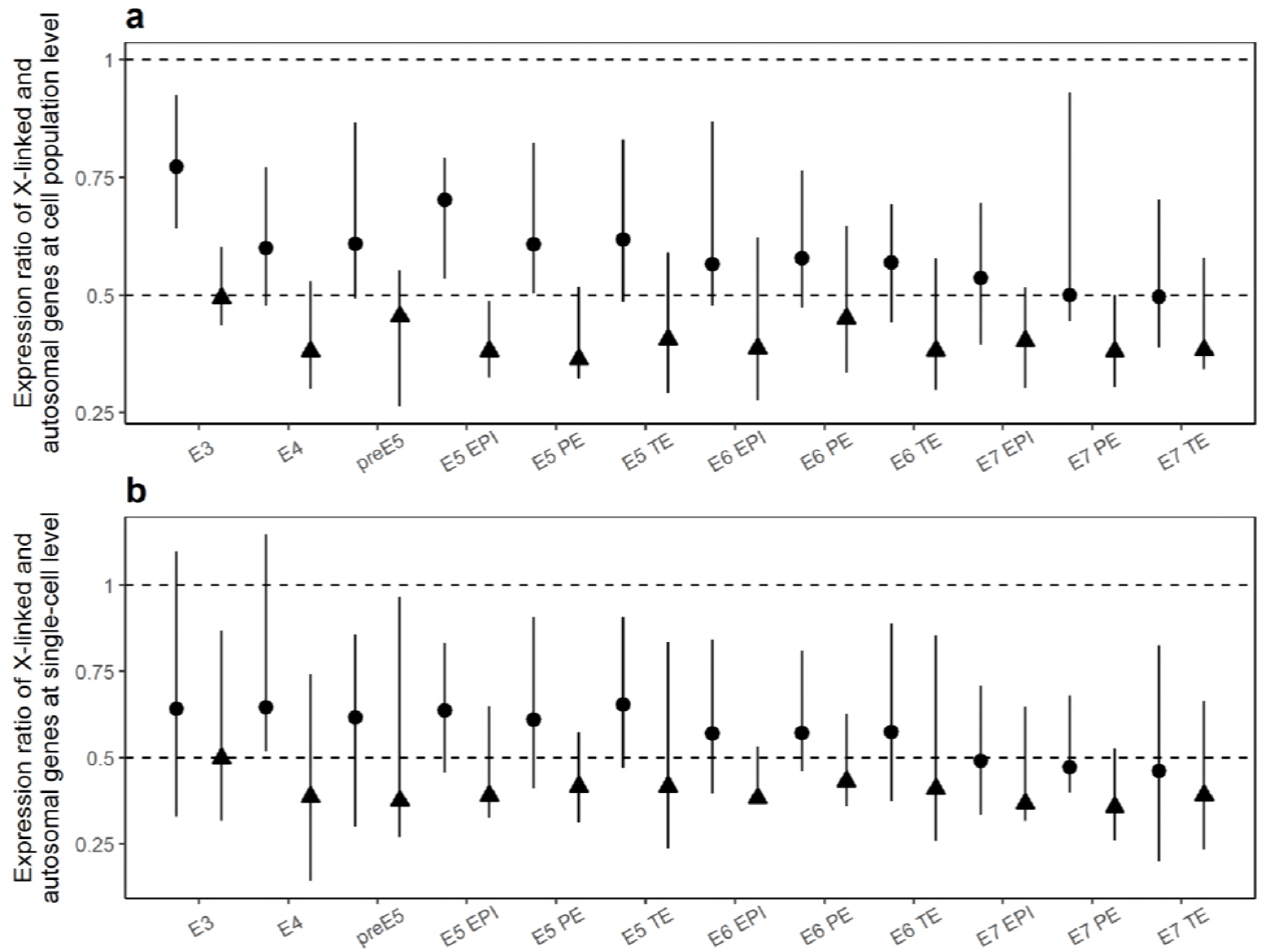
No X-chromosome dosage compensation in human single-cell RNA-seq expression profiles. **(a)** Ratio of the median mRNA expression between X-linked and autosomal genes at the cell population level. Error bars show 90% confidence intervals of the medians, estimated by respectively bootstrapping X-linked and autosomal genes 1000 times. **(b)** Ratio of median mRNA expression between X-linked and autosomal genes at single-cell level. Error bars show 100% confidence intervals of the medians at each cell. In all panels, data from male and female cells are represented by triangles and circles, respectively. Two-tailed Mann–Whitney U test was used to test the equality of the mean expression ratio with 1 (filled symbols, *P*<0.05; open symbols, *P*≥0.05). E3 to E7 indicate embryonic days of the trophectoderm (TE), primitive endoderm (PE) and epiblast (EPI) lineages.

To further assess the dosage imbalance between X chromosome and autosomes beyond the cell population-average expression, we computed the X:AA ratio at the single-cell level. For each cell, we compared the fraction of X-linked genes whose expression level is no less than 10 RPKM (Petropoulos, et al. 2016) and the same fraction of autosomal genes with the highest expression levels in that cell. We then computed the X:AA expression ratio as the ratio of median mRNA expression levels between X-link genes and autosomal genes in each cell. Similar to the population level results, we found that the X:AA expression ratio is ~0.5 in males and gradually decreases to 0.5 in females (**Fig. 1b**). Moreover, the X:AA expression ratio is always below 1 after E4 in all male and female cells, suggesting that the dosage of expression between X chromosome and autosomes is imbalanced in all cells from E5 onward (**Fig. 1b**).

Together with previous observations (Chen and Zhang 2016; Xiong, et al. 2010), these results demonstrate an overall lack of X upregulation at the mRNA level in both male and female preimplantation cells, despite the biallelic expression of X-linked genes in female cells during this period. Furthermore, we found that dosage conversion occurs early, i.e., before the 8-cell stage, and the conversion is fast, such that the X:AA expression ratio reaches ~0.5 in no more than a week.

### The expression noise of X-linked genes is higher than that of autosomal genes in differentiated female cells

The physiological X:AA dosage conversion without interference of normal development implies a lack of phenotypic consequence for different X:AA expression ratios, at least in the range of 0.5 to 1. We thus asked whether X-linked genes are less dosage sensitive than autosomal genes, to which the insensitive X hypothesis would answer “yes”, whereas Ohno’s hypothesis would answer “no”, as it assumes dosage sensitivity for most, especially X-linked genes.

We calculated the coefficient of variation (CV) of mRNA expression for each gene, measured as the standard deviation divided by the mean of single cells with the same lineal status. CV has been considered by some (Kaern, et al. 2005) as a direct and unambiguous measure of expression noise (but see below) compared to the expression differences among biological replicates (Mullon, et al. 2015). We then calculated the ratio of the median CV between X-linked and autosomal genes. For males, this X:AA CV ratio is always larger than 1 regardless of lineage and time point (triangles in **Fig. 2a**). Specifically, the 90% confidence interval of the X:AA CV ratio is always larger than 1 (**Fig. 2a**). This result is consistent with previous theoretical predictions of higher expression noise for haploid-than diploid-expressed genes (Cook, et al. 1998; Wang and Zhang 2011). For females, X-linked genes maintain biallelic expression up to embryonic day 7. Without the lack of ploidy difference, the X:AA CV ratio is not expected to be higher than 1. We found that in female cells, the CV ratio is slightly higher than 1 from E3 to E5, with the 90% confidence interval overlapping with 1. For female cells on E6 and onward, the CV ratio is significantly higher than 1 (circles in **Fig. 2a**), which might be caused by lowered expression of female X-linked genes during this time period (Fig. 1). Despite this confounding factor (see below for a better controlled analysis), these findings are consistent with noisier expression of diploid X-linked genes, and therefore lower dosage sensitivity for X-linked genes than autosomal genes.

**Figure 2.**
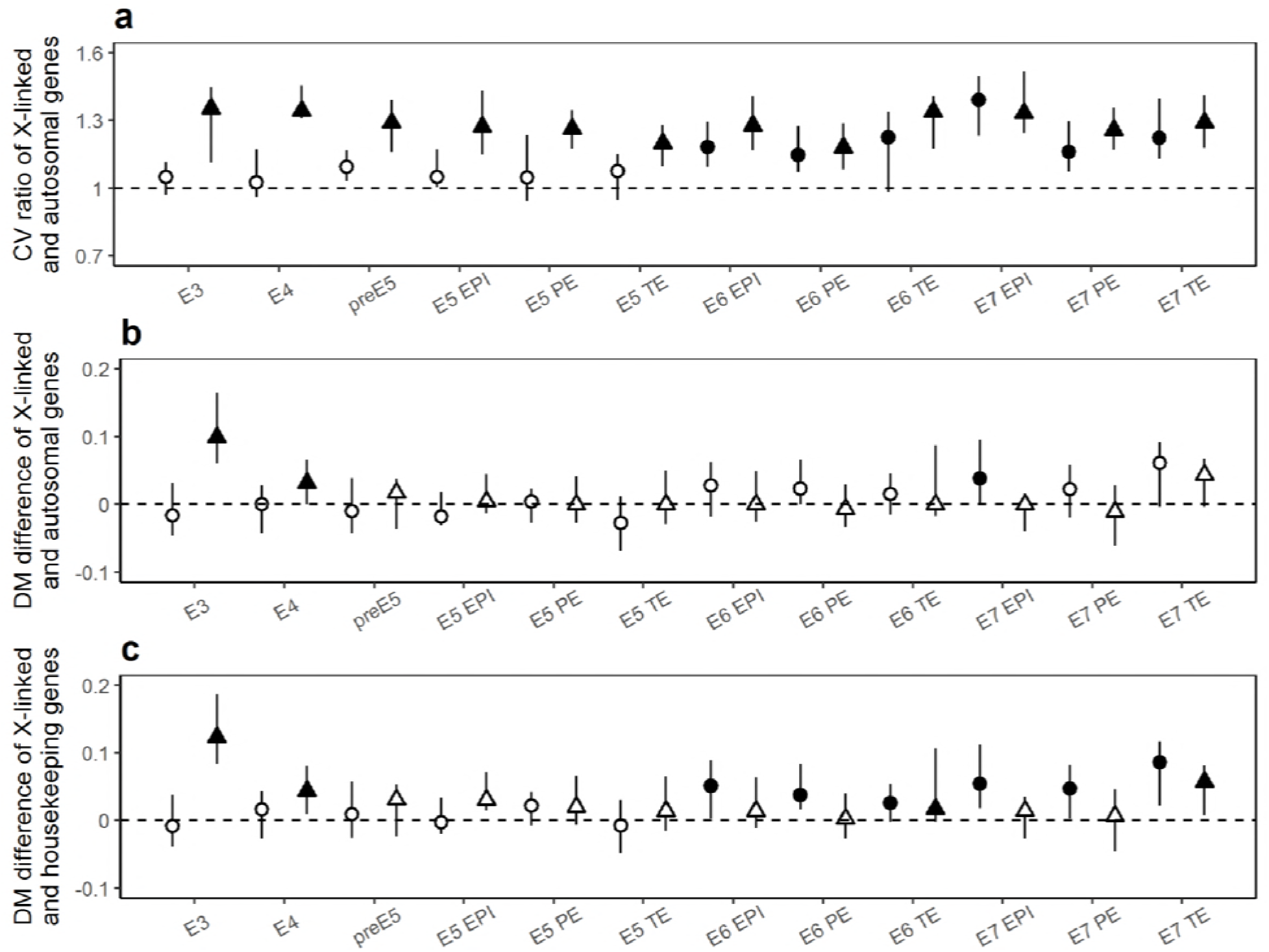
The insensitive X hypothesis is more likely to be true compared to Ohno’s hypothesis. **(a)** Ratio of the median CV between X-linked and autosomal genes. **(b)** Difference between the median DM of X-linked and that of autosomal genes. **(c)** Difference between the median DM of X-linked and that of housekeeping genes. In all panels, data from male and female cells are represented by triangles and circles, respectively. Error bars show the 90% confidence intervals of the medians, estimated by respectively bootstrapping X-linked and autosomal or housekeeping genes 1000 times. Two-tailed Mann–Whitney U test was used to test the equality of the CV ratio with 1 (a) or the DM difference from 0 (b and c) (filled symbols, *P*<0.05; open symbols, *P*≥0.05). E3 to E7 indicate embryonic days of the trophectoderm (TE), primitive endoderm (PE) and epiblast (EPI) lineages. Twenty genes with similar expression levels as the focal gene were used to compute DM.

Because the expression noise represented by CV is confounded by the expression level of the gene, commonly recognized as the finite-number effect (Kaern, et al. 2005), we calculated DM, the ***D***istance of its noise (CV) to the ***M***edian noise (CV) of the genes with comparable mean expression levels (Newman, et al. 2006). Genes with bigger DM are noisier than expected based on the expression level and, therefore, should be less dosage-sensitive. Because DM is defined as the linear distance between the CV of a specific gene and the median CV of genes with similar mean expression levels, DM values should also be compared linearly, i.e., by subtraction, instead of division (as in the case of CV). We found that the median DM of X-linked genes is larger than that of autosomal genes in 5 out of 12 examined male lineages, which was significant in two lineages. In contrast, two male lineages show the opposite trend, though neither is significant (triangles in **Fig. 2b**). These observations favor the insensitive X hypothesis over Ohno’s theory, albeit not significantly (5 *vs* 2 or 2 *vs* 0). Moreover, haploid genes should theoretically be noisier than diploid genes for similar expression levels (Cook, et al. 1998; Wang and Zhang 2011), but we found no significant increase in DM between X-linked genes and autosomal genes after E4 (**Fig. 2b**). The slightly higher DM of X compared to AA in early time points is likely a transition state as the maternal to zygotic transition occurs. Combined with the above result on CV (**Fig. 2a**), these findings suggest that in the male preimplantation cells, the apparent dosage sensitivity of X-linked genes is at least partly due to the finite-number effect, i.e., low expression levels relative to autosomal genes.

On the other hand, in differentiated female lineages from E6 onward, the median DM of X-linked genes is always larger than that of autosomal genes in 6 examined lineages, which is significant for one lineage (circles in **Fig. 2b**). This pattern, which is supportive of the insensitive X hypothesis, remains qualitatively the same when different numbers of genes with comparable mean expression are used to calculate DM (**Fig. S2**). In addition to the comparison of CV, these findings suggest that at least for differentiated female cells, the insensitive X hypothesis, which is not caused by either the finite-number effect of expression level or the ploidy differences between X and autosomes, is more likely to be true than Ohno’s hypothesis.

Notably, this result is inconsistent with a microarray-based study, which claimed that transcriptional variation of X-linked genes is not different from that of autosomal genes both before and after controlling for transcript abundance (Yin, et al. 2009). However, this result could be explained by the inability of microarrays to detect variations at the single-cell level and/or small expression differences among genes (Xiong, et al. 2010).

### Housekeeping genes exhibit less noise and are preferentially located on autosomes

To further assess the dosage sensitivity of X-linked genes, we compared the DM values of X-linked genes with those of housekeeping genes (Eisenberg and Levanon 2013), which are widely considered as dosage-sensitive (Bar-Even, et al. 2006; Newman, et al. 2006). By first confirming the reduced expression noise of housekeeping genes (**Fig. S3**), we compared expression noise of X-linked genes to housekeeping genes. We found that the median DM of X-linked genes is larger than that of housekeeping genes in all twelve male cell lineages, among which four are statistically significant (**Fig. 2c**). This observation is consistent with the expected higher noise of haploid expressed genes (Cook, et al. 1998; Wang and Zhang 2011). On the other hand, female cells always exhibit significantly noisier expression for X-linked genes than housekeeping genes from E6 onward (**Fig. 2c**), suggesting that X-linked genes are less dosage-sensitive than housekeeping genes after controlling for the finite-number effect and ploidy differences.

The haploid expression nature and lack of a general mechanism for dosage balancing with autosomal genes make X an undesirable location for dosage-sensitive genes. The insensitive X hypothesis thus also predicts a depletion of housekeeping genes on X. We found that among one-to-one orthologs in chicken, 53 housekeeping genes are located on the X chromosome (out of 360 X-linked genes), which is proportionally less than 2755 out of 11649 genes on autosomes (*P* = 10^-4^, Chi-squared test). As housekeeping genes are widely expressed in different tissues, this result is consistent with previous observations that the breadth of expression is lowered for X-linked genes (Hurst, et al. 2015; Lercher, et al. 2002).

The depletion of housekeeping genes in X chromosome may have evolved via two scenarios: (i) a chromosome depleted of housekeeping genes becomes a sex chromosome, or (ii) housekeeping genes are removed from the X chromosome once recombination between the therian X and Y is halted. Supporting the latter scenario, a previous study on out-of-X gene movement found that autosomal retrogenes functionally compensate for the silencing of their X-linked housekeeping parental genes (Potrzebowski, et al. 2008). However, dating analyses revealed that retrogenes have been produced since the common ancestor of mammals, whereas the selection for functional compensation driving retrogene export from the X chromosome began much later (Potrzebowski, et al. 2008).

Thus, we tested the other scenario, i.e., whether X chromosome had evolved from an autosome depleted of housekeeping genes. Chicken chromosome 1 and 4 consist of regions syntenic to the human X chromosome (International Chicken Genome Sequencing Consortium 2004). We thus respectively compared the fraction of housekeeping genes among all genes with one-to-one orthologs on chicken chromosome 1 and 4 with that on other chicken autosomes.

Both chromosome 1 (*P* < 10^-4^, Chi-squared Test) and 4 (*P* = 0.002, Chi-squared Test) were found to have significantly lower fractions of housekeeping genes than other autosomes (**Table 1**).

**Table 1.** Chromosomes with lower than average numbers of housekeeping genes are predisposed to become sex chromosomes.

| Chromosome | No. of housekeeping genes with one-to-one orthologs |  | No. of all genes with one-to-one orthologs |  | *P* value<br>(Chi-squared Test) |
| --- | --- | --- | --- | --- | --- |
|  | chicken chr1/4 | other chicken autosomes | chicken chr1/4 | other chicken autosomes |  |
| Complete chromosome |  |  |  |  |  |
| Chicken chr1 | 316 | 2384 | 1615 | 9840 | < 10<sup>-4</sup> |
| Chicken chr4 | 160 | 2540 | 835 | 10620 | 0.002 |
| Syntenic regions |  |  |  |  |  |
| Chicken chr1 | 205 | 2495 | 1065 | 10390 | 0.0006 |
| Chicken chr4 | 149 | 2551 | 802 | 10653 | 0.0006 |

This result remains when only the syntenic (to human X) fractions of chromosome 1 and 4 are analyzed (**Table 1**, see Method). The finding that the human X chromosome evolved from autosomes or part of autosomes depleted of housekeeping genes is in line with selective pressure against X-linkage for dosage-sensitive genes. Collectively, our results suggest that X-linked genes are significantly noisier than well-defined dosage-sensitive genes and generally not as dosage sensitive as autosomal genes.

## Conclusions

Our reanalyses of the single-cell transcriptome data of preimplantation human embryos revealed that male X-linked genes are not two-fold upregulated from the 8-cell stage to the time-point just prior to implantation, during which female X-linked genes gradually decrease their expression from a gametic X:A ~ 1 to a zygotic X:AA ~ 0.5. Consequently, although dosage balance between sexes is achieved before implantation, the expression levels of X-linked genes are not dosage-balanced with autosomal genes. In addition, we found that X-linked genes in differentiated female cells have higher expression noise than autosomal genes. Such an observation contrasts with the primary assumption of Ohno’s hypothesis, i.e., that most X-linked genes are at least as dosage-sensitive as autosomal genes. Moreover, comparative analysis with the chicken genome revealed that X chromosome likely originated from autosomes or part of autosomes that were depleted of housekeeping genes, suggesting selective pressure against X-linkage for dosage-sensitive genes.

We propose here an alternative to Ohno’s hypothesis, i.e., the “insensitive X hypothesis”, to explain the insensitivity of X-linked genes to the two-fold expression change caused by either evolutionary degeneration of Y-linked homologs, X-inactivation or the physiological transition of ploidy during early embryonic development. We utilized recently published single-cell RNA-seq data of human embryos (Petropoulos, et al. 2016) and measured expression noise as a proxy for dosage sensitivity (Mullon, et al. 2015). The biallelic expression (Petropoulos, et al. 2016) of X-linked genes in female cells allows exclusion of noise elevation due to haploid expression (Cook, et al. 1998; Wang and Zhang 2011). Supporting the “insensitive X hypothesis”, our empirical analysis suggests that X-linked genes are noisier than autosomal genes and are less dosage sensitive than housekeeping genes, at least in the differentiated female preimplantation embryo.

Our study includes some caveats that are worth considering. First, the individual cells were categorized into three clearly segregating lineages (TE, PE and EPI) in which pervasive heterogeneity still exists. However, it is highly unlikely that this source of heterogeneity among single cells influences X chromosomes more than autosomes. Second, instead of directly measuring fitness upon suboptimal expression, dosage sensitivity is inferred from gene expression noise. Although there is evidence for reduced expression noise of genes that are sensitive to dosage (Batada and Hurst 2007; Lehner 2008), fitness effects of gene dosage (Keren, et al. 2016) assessed at the genomic scale would be helpful to further test the insensitive-X hypothesis.

How do organisms with incomplete or no dosage compensation avoid deleterious effects of gene dose differences? A previous study in chicken showed that ohnologs, which are duplicated genes known to be dosage-sensitive, are preferentially dosage-compensated on the chicken Z chromosome (Zimmer, et al. 2016). As we showed in this study, X-linked genes exhibit noisier expression, and thus, gene-specific dosage compensation may still be suboptimal for X-linked dosage-sensitive genes, especially on the single-cell level. Therefore, dosage-sensitive genes are preferentially autosomal, which is achievable by two evolutionary scenarios. One possibility is that dosage-sensitive genes had been removed from the X chromosome (Potrzebowski, et al. 2008). Alternatively, we propose here that X chromosome has evolved from an ancestral autosome that was depleted of dosage-sensitive genes. This latter scenario is supported by comparison between the human X chromosome and the chicken genome. Because selection-driven gene export from the X chromosome began after the recombination between the therian X and Y was halted (Potrzebowski, et al. 2008), evolution of X from chromosomes with fewer dosage-sensitive genes is an evolutionary trajectory with a lower fitness cost for the intermediate genotypes.

In the future, it would be interesting to generate single-cell proteomic data from human cells to validate the above findings at the proteomic level, as was recently carried out for mean protein abundance of a human diploid cell population (Chen and Zhang 2015). It would also be interesting to confirm our results by comparing human haploid transcriptomic or proteomic data with the corresponding data from a bird, as previously reported for diploid transcriptomic data (Julien, et al. 2012; Lin, et al. 2012).

## Materials and Methods

Gene models and mapping of EnsEMBL gene IDs to UniProt/SwissProt accessions in human were downloaded from EnsEMBL (release 87) (Cunningham, et al. 2015). Human and chicken one-to-one orthologs were also downloaded from the same release of EnsEMBL. Synthetic regions of human X chromosome in the chicken chromosome 1 and 4 were previously identified (International Chicken Genome Sequencing Consortium 2004) and further constrained here to between the first and last one-to-one ortholog within that region. Genes expressed uniformly across a panel of tissues captured by RNA-seq are identified as human housekeeping genes (Eisenberg and Levanon 2013). The number of RNA-seq reads per kilobase of exon per million reads mapped (RPKM), supplied by the original authors of the study, was used to measure gene expression levels (Petropoulos, et al. 2016). To avoid the effect of technical noise in single-cell expression measurements, especially for lowly expressed genes, we followed previous procedures and focused on X-linked genes with RPKM ≥10 (Petropoulos, et al. 2016). We also tried a less stringent cut-off (RPKM ≥5). At the cell population level and for each embryonic day and each lineage, we then compared the fraction of X-linked genes whose expression level was at least 10 or 5 RPKM with the same fraction of autosomal genes that had the highest expression level (Chen and Zhang 2016). In addition, at the single-cell level for each cell, we compared the fraction of X-linked genes whose expression level was at least 10 or 5 RPKM with the same fraction of autosomal genes that had the highest expression level. We then computed the ratio of the median mRNA expression level between X-link genes and autosomal genes. To compare expression noise, we calculated the CV of mRNA expression in each gene at the cell population level, measured as the standard deviation divided by the mean, and computed the ratio of the median CV between X-link genes and autosomal genes. As another measurement of expression noise, we used DM, which was calculated as follows. We ranked the genes by their mean expressions, and then for each specific gene, we used 10, 20 or 50 genes with similar levels of focal gene to calculate the median CV, and the difference in the median CV and the focal CV was used as the DM (Newman, et al. 2006). We then calculated the difference in median DM between X-link genes and autosomal genes, as well as between X-link genes and housekeeping genes.

## Acknowledgements

We thank Jianzhi Zhang for valuable comments. This work was supported by Project 2017YFA0103504 of the National Key R&D Program of China awarded to X.C, and the National Natural Science Foundation of China Project 31671320 awarded to J.-R.Y and Project 31771406 awarded to X.C.

## Figure legends

**Figure S1.**
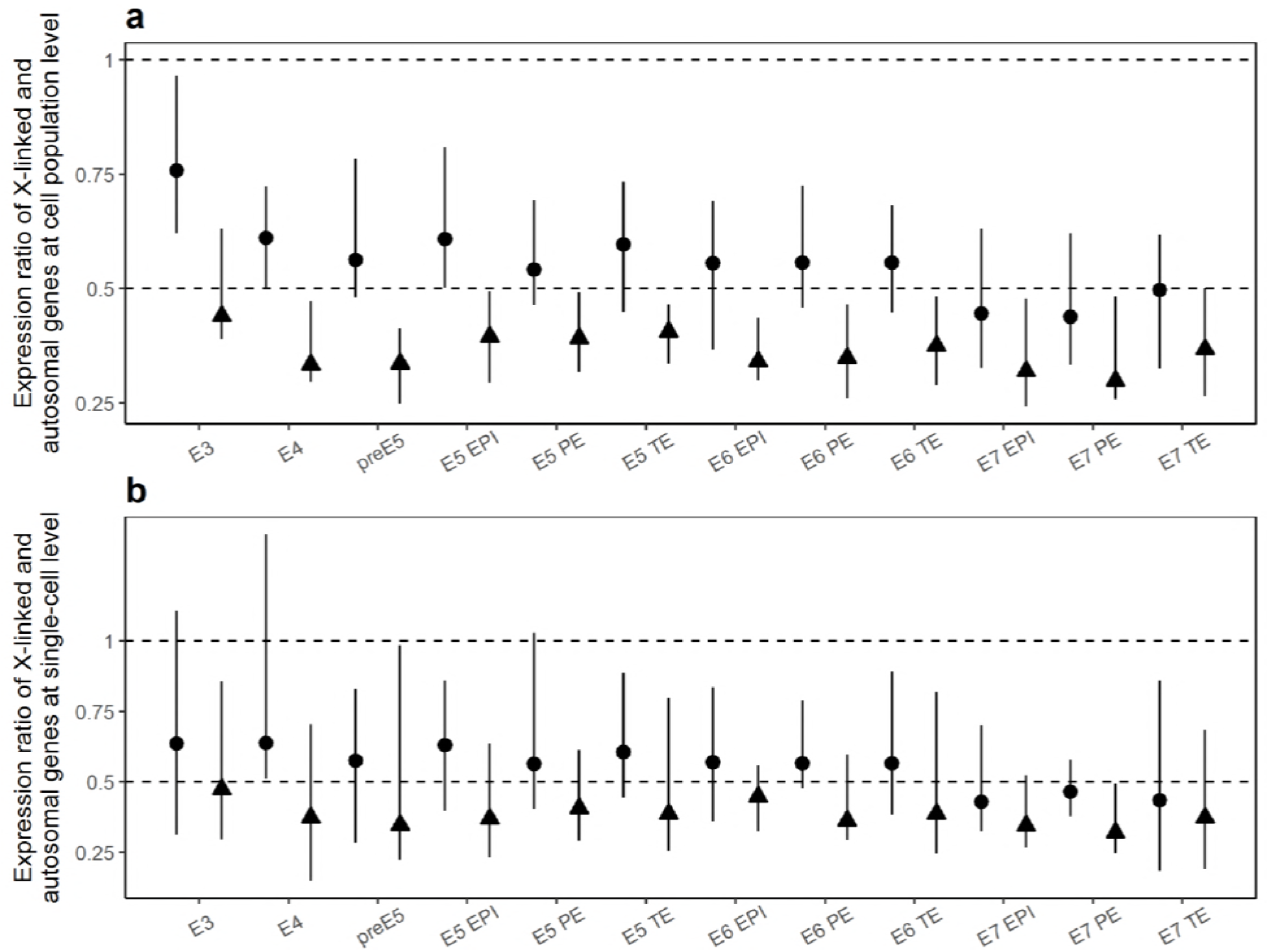
No X-chromosome dosage compensation in human single-cell RNA-seq profiling.

Similar to Fig. 1 except that X-linked genes with RPKM no less than 5 are considered.

**Figure S2.**
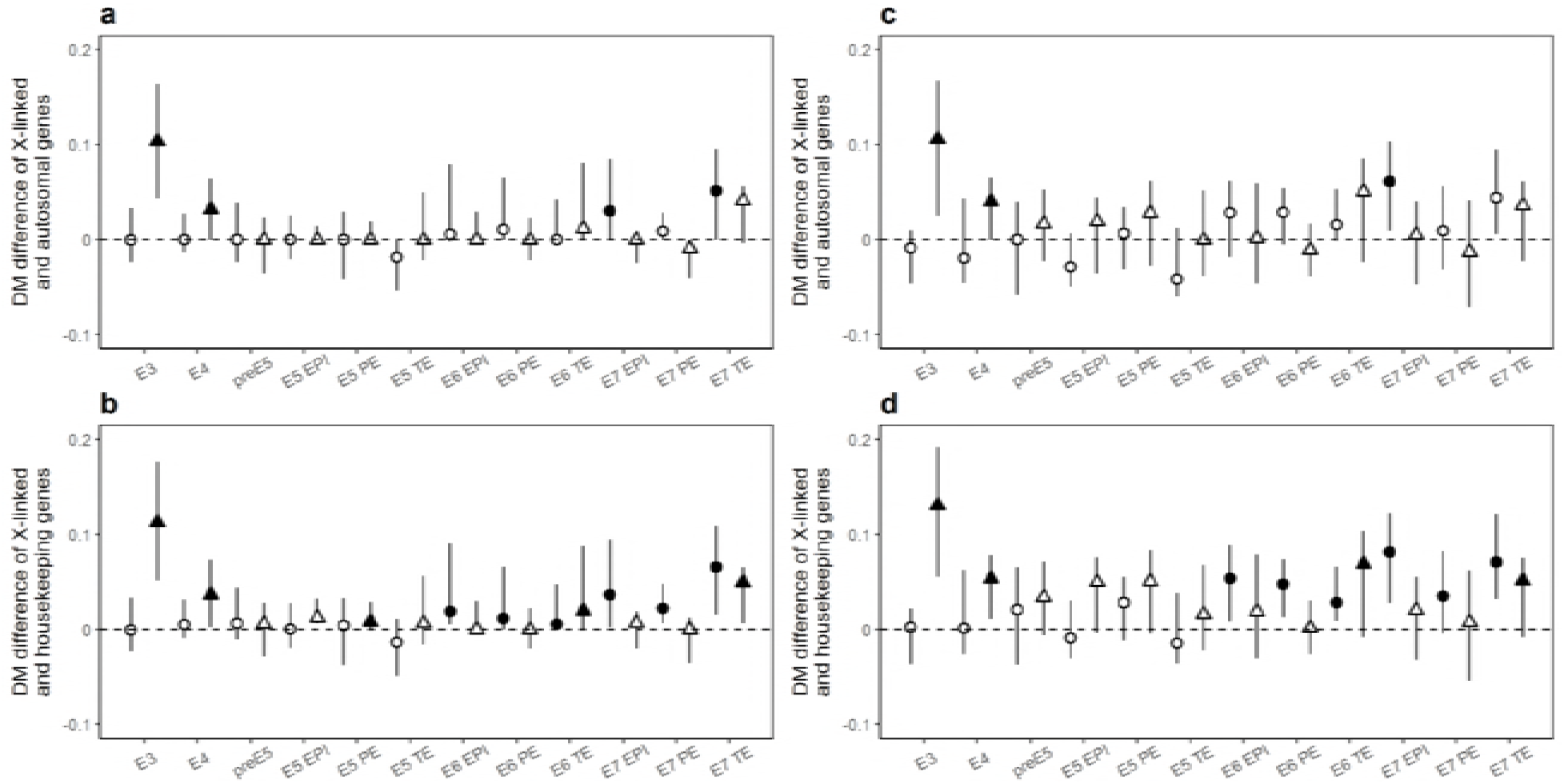
The insensitive X hypothesis is more likely to be true than Ohno’s hypothesis. **(a and b)** Similar to Fig. 2**b and c** except that 10 genes with similar expression levels as the focal gene are used to compute DM. **(c and d)** Similar to Fig. 2**b and c** except that 50 genes with similar expression levels as the focal gene are used to compute DM.

**Figure S3.**
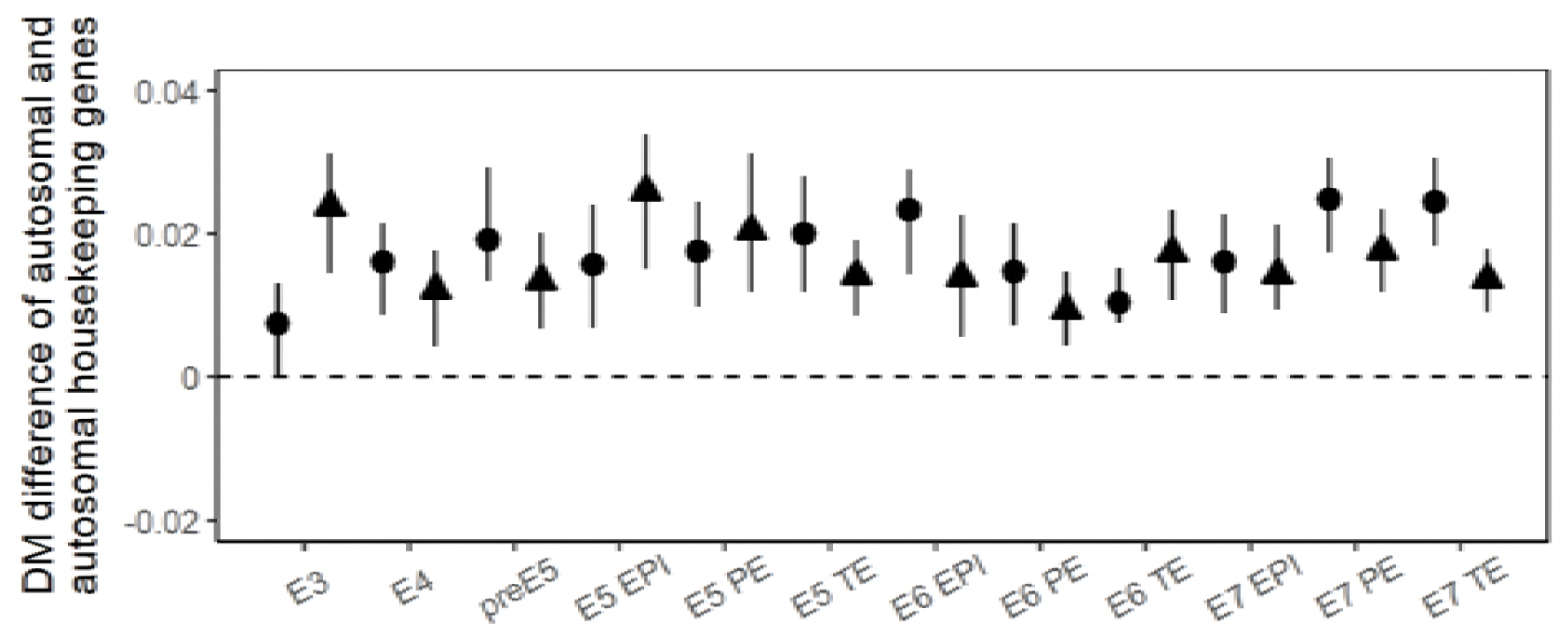
Housekeeping genes are more dosage sensitive than other autosomal genes. Similar to Fig. 2b except that the DM of autosomal housekeeping genes is compared to that of autosomal genes.

